# Reduced Resting-State Connectivity in the Precuneus is correlated with Apathy in Patients with Schizophrenia

**DOI:** 10.1101/489161

**Authors:** Caroline Garcia Forlim, Leonie Klock, Johanna Bächle, Laura Stoll, Patrick Giemsa, Marie Fuchs, Nikola Schoofs, Christiane Montag, Jürgen Gallinat, Simone Kühn

**Author notes:** indicates a shared first-authorship (Caroline Garcia Forlim, Leonie Klock).

## Abstract

A diagnosis of schizophrenia is associated with a heterogeneous psychopathology including positive and negative symptoms. The disconnection hypothesis, an early pathophysiological framework conceptualizes the diversity of symptoms as a result from disconnections in neural networks. In line with this hypothesis, previous neuroimaging studies of patients with schizophrenia reported alterations within the default mode network (DMN), the most prominent network at rest.

Aim of the present study was to investigate the functional connectivity during rest in patients with schizophrenia and healthy individuals and explore whether observed functional alterations are related to the psychopathology of patients. Therefore, functional magnetic resonance images at rest were recorded of 35 patients with schizophrenia and 41 healthy individuals. Independent component analysis (ICA) was used to extract resting state networks.

Comparing ICA results between groups indicated alterations only within the network of the DMN. More explicitly, reduced connectivity in the precuneus was observed in patients with schizophrenia compared to healthy controls. Connectivity in this area was negatively correlated with the severity of negative symptoms, more specifically with the domain of apathy.

Taken together, the current results provide further evidence for a role DMN alterations might play in schizophrenia and especially in negative symptom such as apathy.

## 1. Introduction

The diagnostic category of schizophrenia summarizes heterogeneous symptoms that range from so-called positive symptoms, which refer to an excess of function with prominent examples such as hallucinations and delusions, to so-called negative symptoms, which refer to a diminishment of function such as anhedonia, affective blunting, and apathy (Andreasen, 1985). Furthermore, a key feature of schizophrenia is an alteration in the subjective experience of a continuous sense of self, which has been argued to account for both positive and negative symptoms (Sass and Parnas, 2003). Regarding the pathophysiology of schizophrenia, an early and influential hypothesis proposes that core symptoms of schizophrenia can be regarded as dysfunctional interactions between different brain regions rather than pathological changes within localized areas (Friston et al., 2016; Friston and Frith, 1995). Furthermore, it has been suggested that these alterations in the connectivity of neural circuits can account for the heterogeneity of symptoms that are associated with a diagnosis of schizophrenia as they are thought to affect the coordination of mental processes (Andreasen et al., 1998)

Interactions in the brain can be described by means of complex networks. A neural network, when applied to functional magnetic resonance imaging (fMRI), is composed of cortical areas that are functionally connected with each other. Functional connections describe temporal correlations of distributed neural areas. Brain networks are commonly computed on data of individuals who were lying without a specific task instruction in an MRI scanner, a so-called resting-state (rs-fMRI). Brain networks at rest represent intrinsic or “default” brain activity and therefore, are of fundamental importance for the understanding of the human brain (Raichle et al., 2001). Although fMRI recordings during the performance of a cognitive task (task-fMRI) are widely used and accepted as a valid tool to investigate neural correlates of specific behaviours, the energy consumption during a task is associated with an increase of less than 5% from the brain’s baseline energy consumption (Raichle and Mintun, 2006) - indicating that the brain is constantly active at rest. Importantly, task and resting state fMRI can be understood as complimentary rather than opposing modalities that are focusing on different theoretical and technical aspects: task-fMRI investigates cortical activation sites and rs-fMRI focuses on temporal aspects and correlations among sites to build brain networks. Rs-fMRI may be a promising method to study the pathophysiology of schizophrenia as it is comparatively easy to acquire in a clinical population, does not include differences in task performance that might potentially confound group fMRI analyses (Whitfield-Gabrieli and Ford, 2012a), and has been related to a fundamental aspect of human experience, the sense of self (Raichle and Raichle, 2001).

A reliable method of extracting the brain’s resting state networks is a technique borrowed from engineering called independent component analysis (ICA). This method blindly recovers source signals from a mixture of sources (McKeown et al., 2003) and can be illustrated by means of the classic cocktail party problem: microphone recordings that contain simultaneous conversations of people can be used to recover individual voices of people by employing the method of ICA. The same principle can be applied to brain signals in rsfMRI - ICA decomposes the brain’s activity into multiple sources (components). One advantage of this method is that it is data-driven and requires no prior assumptions as is the case for another popular method called seed-based functional connectivity (Fox and Raichle, 2007). Each component (source) that is retrieved by ICA is a spatial grouping of voxels with temporally coherent activity. Depending on the spatial grouping of voxels, the components are associated with sources that are either related to brain activity or to noise such as movement, blinking, breathing, and heartbeat. The main brain activity-related sources that can be retrieved with ICA “closely resemble discrete cortical functional networks” (Beckmann et al., 2005), and are named resting state networks (RSN): default mode (DMN), basal ganglia, auditory, visuospatial, sensory-motor, salience, executive control, language and visual networks.

The term “default mode” refers to a set of brain regions that exhibit increased and coherent neural activation during resting conditions compared to conditions requiring an external attention focus. As it is conceptualized as the baseline of brain functioning at rest, the DMN became the most studied RSN comprising key cortical regions such as the posterior cingulate cortex, precuneus, medial prefrontal cortex, hippocampus, insula, and inferior parietal cortices. Interestingly, studies investigating the DMN of patients diagnosed with schizophrenia have found functional alterations in this RSN (Calhoun et al., 2009; Karbasforoushan and Woodward, 2012; Whitfield-Gabrieli and Ford, 2012b). More precisely, studies that employed ICA to investigate the DMN showed connectivity in various brain areas to be either decreased (Camchong et al., n.d.) (Rotarska-Jagiela et al., 2010) (Orliac et al., 2013), increased (Garrity et al., 2007), or both (Mannell et al., 2010) (Li et al., 2017) (Ongür et al., 2010). However, other studies did not report any differences in the DMN between patients with schizophrenia and healthy individuals (Wolf, 2011). As the results of these previous ICA studies remain inconclusive, we set out in the current study to explore functional connectivity in resting-state networks using ICA in a larger group of patients diagnosed with schizophrenia compared to matched healthy individuals and whether these alterations are related to the severity of clinical symptoms of schizophrenia patients. Based on the reviewed literature we hypothesized connectivity alterations in the DMN.

## 2. Methods

### 2.1 Participants

In total, 76 participants were included in the reported analysis of which 41 were healthy individuals and 35 individuals met the criteria for a diagnosis of schizophrenia following the International Classification for Diseases and Related Health Problems (ICD-10) (World Health Organization, 2012). Patients diagnosed with schizophrenia were recruited at St. Hedwig Hospital, Department for Psychiatry and Psychotherapy of the Charité-Universitätsmedizin Berlin (Germany). Healthy individuals were recruited using online advertisements and flyers and did not meet the criteria for any psychiatric disorder based on information acquired with the Mini International Neuropsychiatric Interview (MINI) (Ackenheil et al., 1999) and were not in current or past psychotherapy of an ongoing mental health-related problem. MRI exclusion criteria such as claustrophobia, neurological disorders and metallic implants applied to all participants. Healthy individuals matched the group of patients in terms of age, sex, handedness and level of education (Table 1). Handedness was acquired with Edinburgh Handedness Inventory (Oldfield, 1971) (*n*=75), cognitive functioning was tested using the Brief Assessment of Cognition in Schizophrenia (Keefe et al., 2008) (*n*=65) and verbal intelligence with a German Vocabulary Test (Schmidt and Metzler, 1992) (*n*=72). Supplemental information provides details regarding the medication of patients. All procedures of the study were approved by the ethics committee of the Charité-Universitätsmedizin Berlin.

**Table 1.** Sample Description.

|  | **Healthy**<br>**Participants** | **Schizophrenia**<br>**Patients** | **Statistics** |  |
| --- | --- | --- | --- | --- |
|  | Mean (*SD*) | Mean (*SD*) | T (*DF*) | P Value |
| **Sociodemographic Characteristics** |  |  |  |  |
| Number of participants | 41 | 35 |  |  |
| Age (years) | 35.2 (*11.0*) | 35.3 (*10.8*) | -0.059 (*74*) | 0.953 |
| Gender | 24 male / 17 female | 21 male / 14 female |  |  |
| Edinburg Handedness Inventory<sup>a</sup> | 79.4 (*38.5*) | 75.56 (*52.6*) | 0.331 (*59*) | 0.742 |
| Education (years) | 14.1 (*2.9*) | 13.1 (*3.8*) | 1.345 (*74*) | 0.183 |
| BACS<sup>a</sup> | 270.4 (*37.0*) | 234.1 (*28.4*) | 4.427 (*63*) | < 0.001 |
| Verbal Intelligence (IQ)<sup>a</sup> | 100.5 (*10.6*) | 94.4 (*12.8*) | 2.204 (*70*) | 0.031 |
| **Psychopathology** |  |  |  |  |
| Illness duration (years) |  | 9.4 (*8.8*) |  |  |
| Illness onset (age in years) |  | 25.6 (*8.9*) |  |  |
| Chlorpromazine-equivalent (mg) |  | 317.1 (*221.6*) |  |  |
| SANS Composite Score<sup>a</sup> |  | 20.2 (*12.0*) |  |  |
| SAPS Composite Score<sup>a</sup> |  | 15.4 (*14.9*) |  |  |

SD, standard deviation; BACS, Brief Assessment of Cognition in Schizophrenia; SAPS, Scale for Assessment of Positive Symptoms; SANS, Scale for Assessment of Negative Symptoms;
<sup>a</sup> sum score of items reported.

### 2.2 Assessment of Psychopathology

Trained clinicians rated the severity of symptoms with the Scale for Assessment of Negative Symptoms (SANS) (Andreasen, 1989) and the Scale for Assessment of Positive Symptoms (SAPS) (Andreasen, 1984). Table 1 includes details of patients’ psychopathology. The SANS assesses negative symptoms within the domains of affective blunting, alogia, avolition-apathy, anhedonia-asociality, attentional impairment and the SAPS assesses positive symptoms in the domains of hallucinations, delusions, bizarre behavior, and positive formal thought disorder. Both scales rate severity of symptoms on a scale from 0 (absent) to 5 (severe). For both scales a composite score of all items was computed. Additionally, both scales contain a global assessment item of each subdomain.

### 2.3 MRI Data acquisition

Images were collected on a Siemens Tim Trio 3T scanner (Erlangen, Germany) using a 12-channel head coil. Structural images were obtained using a T1-weighted magnetization prepared gradient-echo sequence (MPRAGE) based on the ADNI protocol (TR=2500ms; TE=4.77ms; TI=1100ms, acquisition matrix=256×256×176; flip angle = 7°; 1×1×1mm3 voxel size). Whole brain functional resting state images during 5 minutes were collected using a T2*-weighted EPI sequence sensitive to BOLD contrast (TR=2000ms, TE=30ms, image matrix=64×64, FOV=216mm, flip angle=80°, slice thickness=3.0mm, distance factor=20%, voxel size 3×3×3mm3, 36 axial slices). Before resting state data acquisition was started, participants were in the scanner for about 10 minutes during which a localizer and the anatomical images were acquired so that subjects could get used to the scanner noise. During resting state data acquisition participants were asked to close their eyes and relax.

### 2.4 Preprocessing of resting state data

To ensure for steady-state longitudinal magnetization, the first 5 images were discarded. The acquired data was corrected for slice timing and realigned. Structural individual T1 images were coregistered to functional images and segmented into gray matter, white matter, and cerebrospinal fluid. Data was then spatially normalized to the MNI template and spatially smoothed with a 6-mm FWHM to improve signal-to-noise ratio. All steps of data preprocessing were performed using SPM12. In addition, to control for motion, the voxel-specific mean frame-wise displacement (FD; (Power et al., 2012). FD values were below the default threshold of 0.5 for control and patient group (0.15±0.02 and 0.17±0.02, t-test *p*=0.42).

### 2.5 Independent component analysis (ICA)

ICA is a data-driven analysis tool in which source signals are blindly recovered (McKeown et al., 2003) from mixtures of sources. ICA was calculated using GIFT software ((http://icatb.sourceforge.net/; (Calhoun, 2004) in Matlab 2012b using Infomax algorithm to estimate independent sources. ICA was run 20 times and the results were clustered by ICASSO (min cluster size of 16 and max of 20). Preprocessed data from all subjects in both groups were used. The optimal number of spatially independent resting-state networks (*N*) to be extracted was estimated by the software (*N*=21). The networks were identified using predefined templates in GIFT and a-posteriori by experts C.G.F and S.K. Among the ICA components, the default mode network was identified and taken to the second level analysis in SPM12 using a mask for individual networks provided by http://findlab.stanford.edu/functional_ROIs.html. To make use of the full potentiality of the method, we conducted an exploratory analysis with the remaining networks that were concomitantly extracted by ICA: basal ganglia, visual, salience, auditory, executive control, and visuospatial.

Differences between groups were calculated using a two-sample t-test (*p*=0.001 uncorrected), significant threshold was set to *p*<0.05 corrected for multiple comparison (FWE). Mean FD (Power et al., 2012), gender and age were used as covariates. In those clusters where we found significant group differences, we extracted the mean absolute ICA values for each subjected for a posteriori correlation analysis of connectivity with psychopathology.

### 2.6 Correlation with psychopathology

To investigate whether these differences in resting state networks between healthy individuals and patients with schizophrenia were related to the psychopathology of patients diagnosed with schizophrenia, Spearman’s correlation coefficients between resting state network connectivity and severity of symptoms were calculated. The resting state network connectivity was extracted from the cluster in the DMN cluster in which we found group differences: for each subject we extracted the mean absolute ICA value. In a first step, we correlated the connectivity in the DMN cluster with the composite scores of the SANS and SAPS. In case of a significant correlation, we conducted further exploratory correlations with the global ratings of the associated symptom domains. Statistical analysis was performed using SPSS 22. For the correlation with the SAPS and SANS composite scores, a Bonferroni adjusted significance level of 0.025 (0.05/2) was applied. For the correlations with subdomains we used a Bonferroni corrected alpha value of 0.0125 (0.05/4) for the SAPS and of 0.01 (0.05/5) for the SANS.

## 3. Results

### 3.1 Independent component analysis (ICA)

#### 3.1.1 Default mode network analysis

From 21 components chosen automatically in GIFT, the DMN was identified by experts C.G.F and S.K and taken to 2nd level analysis in SPM 12 (*p*<0.001 uncorrected) using masks for individual networks provided by http://findlab.stanford.edu/functional_ROIs.html. The results indicated a significant decrease in the connectivity within the precuneus in patients with schizophrenia compared to healthy individuals (MNI coordinates= 4 -60 36, *T* = 4.23, *p_peak-level FWE corrected_* = 0.037; clustersize (in voxels) = 35; Figure 1A).

**Figure 1.**
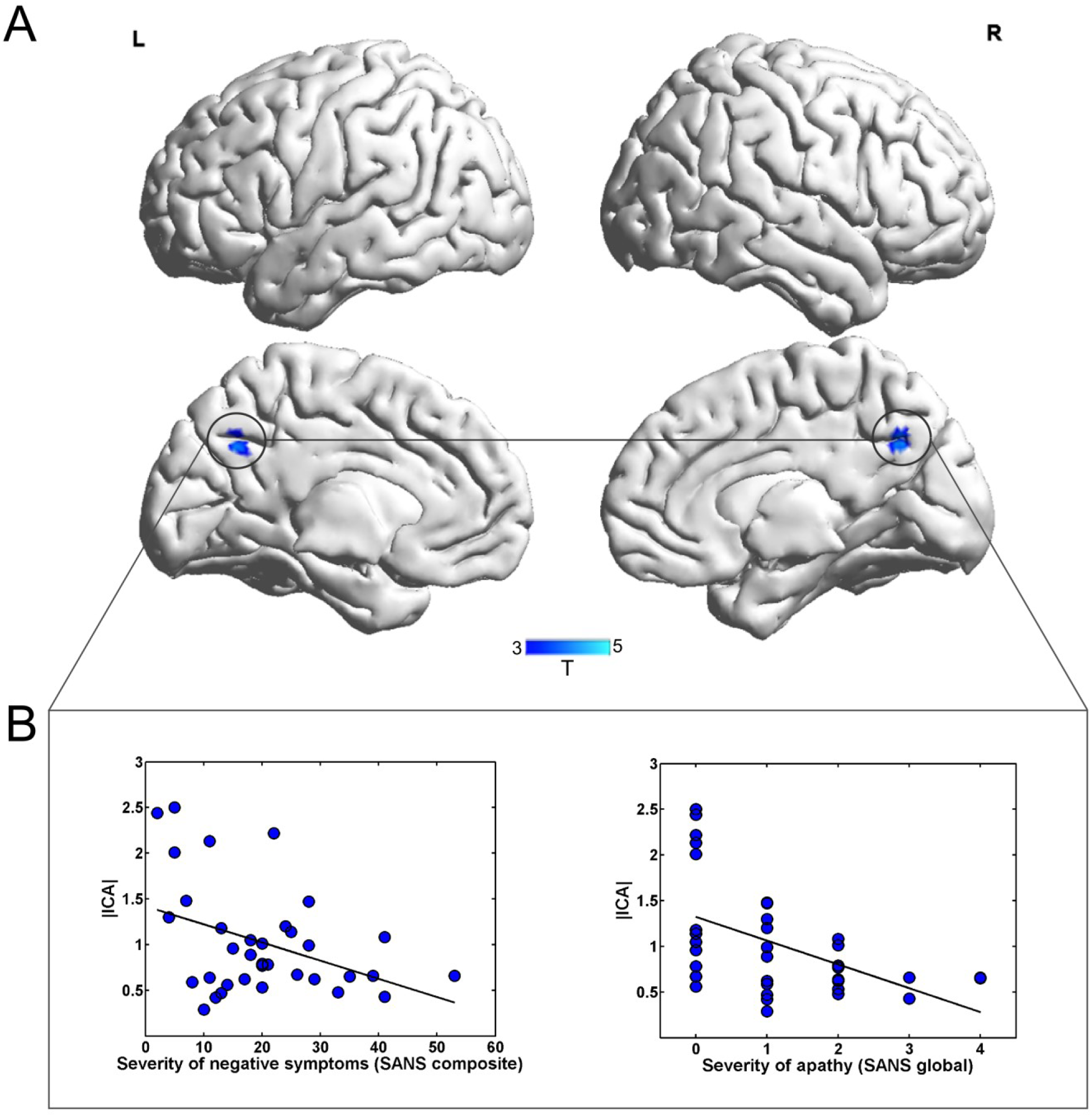
Group differences between patients with schizophrenia and healthy individuals in the DMN. **A.** Group comparison revealed decreased connectivity in the precuneus, as part of the DMN in patients with schizophrenia. **B.** Functional connectivity during rest in the precuneus was significantly related to the severity of negative symptoms (assessed with the SANS) and more specifically with the severity of symptoms regarding avolition/apathy.

#### 3.1.2 Exploratory analysis of the remaining resting state networks

Besides the DMN, ICA method allows to uncover other resting state networks. In order to explore the full potentiality of ICA, we performed further exploratory analysis in additional networks extracted from our dataset. From the total of 21 components automatically estimated, additionally to the DMN, we identified, Basal Ganglia, Visual, Salience, Auditory, Executive control and Visuospatial networks. Following the same procedure for the DMN, the networks were taken to the second level analysis in SPM12 using masks for individual networks provided by http://findlab.stanford.edu/functional_ROIs.html. No significant group differences were found.

### 3.2 Correlational analysis with psychopathology

In order to investigate whether connectivity values of the precuneus cluster in the DMN, given by the mean absolute ICA, were related to psychopathology of patients with schizophrenia, absolute ICA values of the precuneus cluster were extracted and correlated with the composite score of the SAPS (*r*(34) = 0.106, *p* = 0.549) and of the SANS (*r*(35) = -0.398, *p* = 0.018, Figure 1B). Based on the significant relationship between negative symptoms of the SANS and functional connectivity of the precuneus, we explored in a next step whether a specific domain of negative symptoms was associated with the functional connectivity in the precuneus. We observed a significant negative relationship between ICA parameters of the precuneus and avolition-apathy (apathy: *r*(35) = -0.504, *p* = 0.002, Figure 2B) but not any of the other domains (anhedonia *r*(35) = -0.314, *p* = 0.066, affective blunting *r*(35) = -0.165, *p* = 0.343; alogie *r*(35) = -0.235, *p* = 0.175; attention *r*(35) = 0.239, *p* = 0.167).

To further explore the association between apathy and precuneus, we also calculated a multidimensional apathy score following the suggestion of Bortolon and colleagues. (Bortolon et al., 2017). Besides the items assessing behavioral aspects of apathy such as “grooming and hygiene”, “impersistence at school or work”, “physical anergia” this score also includes the items “recreational interests and activities” and “affective nonresponsivity” to account for cognitive and emotional aspects of apathy. That multidimensional apathy score was also significantly correlated with the precuneus (*r*(35) = -0.469, *p* = 0.004).

## 4. Discussion

The DMN network can be considered the baseline of brain processing at rest (Greicius et al., 2003; Raichle et al., 2001) and is assumed to play a crucial role in schizophrenia (Calhoun et al., 2009; Karbasforoushan and Woodward, 2012; Whitfield-Gabrieli and Ford, 2012b). In the presented study, the DMN was extracted from a group of patients diagnosed with schizophrenia and matched healthy individuals using ICA, a powerful data-driven method to infer connectivity that does not require an *a priori* definition of regions of interest. In addition, an exploratory analysis was conducted with all remaining resting state networks concomitantly extracted by ICA. Significant group differences were found solely in the DMN. The group comparison results indicate significantly reduced functional connectivity in the DMN in patients diagnosed with schizophrenia. This finding supplements the results of previous ICA studies that reported heterogeneous findings in multiple brain regions within the DMN: decreased functional connectivity in the posterior cingulate and hippocampus (Rotarska-Jagiela et al., 2010), right paracingulate cortex (Orliac et al., 2013), middle frontal gyrus and anterior cingulate gyrus (Camchong et al., n.d.) and increased functional connectivity in frontal, anterior cingulate and parahippocampal gyri (Garrity et al., 2007). Furthermore, studies observed both decreases and increases (Ongür et al., 2010) as well as no alterations in the DMN (Wolf, 2011).

More specifically, we observed reduced functional connectivity in the precuneus in patients with schizophrenia. Again, previous studies reported both decreased functional connectivity (Li et al., 2017) and increased functional connectivity (Mannell et al., 2010) within the precuneus. Interestingly, the precuneus functions as a central hub within the DMN (Utevsky et al., 2014). Furthermore, we observed that alterations in the functional connectivity of the precuneus in the DMN were negatively related to the severity of negative symptoms. More specifically, we found the avolition-apathy domain to be negatively related to precuneus connectivity in patients. Apathy is a multidimensional syndrome that presents itself as the most common negative symptom of schizophrenia (Bortolon et al., 2017) and is also present in other neuropsychiatric diseases and in Alzheimer’s disease (Mulin et al., 2011; Robert et al., 2009). Apathy is described as a disorder of motivation with reduced or loss of goal-directed behavior, goal-directed cognitive processes, and emotion (Mulin et al., 2011; Robert et al., 2009). Whereas Andreasen (1989) refers to apathy as a decline in energy and drive and thus mainly covers behavioral aspects of apathy in the SANS assessment, Bortolon et al. (Bortolon et al., 2017) emphasize the multidimensional aspects of apathy. Following the proposal of Bortolon et al. (Bortolon et al., 2017) as well as the diagnostic domains proposed by Robert et al. (2009), we calculated in addition to the SANS avolition-apathy score a second apathy score that allows a multidimensional assessment of apathy (see methods). Of note, also the multidimensional apathy score was negatively correlated with the functional connectivity in the precuneus during rest-state further supporting the observed relationship. In sum, the present findings show that patients diagnosed with schizophrenia exhibited reduced functional connectivity in the precuneus of the DMN and that this dysconnectivity was more pronounced in patients with more severe symptoms of apathy, a so-called negative symptom of schizophrenia.

To our knowledge, we are the first to observe a relationship between reduced functional connectivity in the precuneus and apathy. The precuneus has repeatedly been observed to play a central role in remembering events of one’s personal past, referred to as autobiographical memory as part of the episodic memory system that is linked to the storage and recollection of events that contain a strong self-reference (Cabeza and St Jacques, 2007; Cavanna and Trimble, 2006). However, the precuneus is not only involved in remembering one’s past but also in imagining one’s future (Schacter et al., 2007). Also during episodes of rest when the precuneus becomes activated as a functional hub of the DMN, the human mind engages in self-generated thoughts that relate to past and future events (Andrews-Hanna et al., 2014, 2010). Interestingly, on a behavioral level, previous studies report impairments in autobiographical memory in patients diagnosed with schizophrenia (Danion et al., 2005; Riutort et al., 2003; Wood et al., 2006) as well as in generating personal and specific future events (D’Argembeau et al., 2008; de Oliveira et al., 2009; Raffard et al., 2013). Especially deficits in imagining pleasant future events were observed to be associated with apathy symptoms in individuals with schizophrenia (Raffard et al., 2013). According to the authors (Raffard et al., 2013), impairments in the capacity to generate future projection of one’s behavior, especially in preferable future situations, might diminish the motivation to engage in goal-directed behavior as commonly observed in apathy. We speculate that the current results indicating dysconnectivity in the precuneus, which is commonly engaged in future-oriented thoughts, and is related to the severity of apathy provide support from a neural perspective for the observation that impairments in future projections might be related to reduced goal-directed behavior characteristic for apathy.

However, it is not possible to draw causal inferences based on the reported result as the directionality of this relationship still remains unclear. It is also conceivable that the impoverished life circumstances of patients might reduce their motivation to engage in future simulations (de Oliveira et al., 2009) or that diminished goal-directed behavior and cognition, which are characteristic for apathy, might diminish future-oriented thoughts(Raffard et al., 2013), which are potentially associated with the observed precuneus dysconnectivity during resting-state. Future studies are needed to further investigate the direction of this relationship.

### Limitations of the study

A first limitation concerns the fact that 31 of the 35 patients were taking antipsychotic medication. As antipsychotic treatment was shown to affect functional connectivity in patients with schizophrenia (Lui et al., 2010), we addressed this potential confound by calculating and controlling the performed correlation analysis for Chlorpromazine-equivalent (CPZ) (Gardner et al., 2010) (see supplements). Despite the fact that the results remained significant when controlling for CPZ, it would be beneficial for future studies to include antipsychotic-naive patients to exclude medication as a potential confound. A second limitation is that post-hoc correlational analysis with different psychopathology scales might have increased Type I error rates. Therefore, we applied Bonferroni correction for all correlational analysis. A further limitation is that we cannot infer a causal direction of the observed DMN dysconnectivity results emphasizing the need of future longitudinal studies that investigate functional connectivity during rest over the course of illness. Finally, the number of components that is chosen in ICA is an arbitrary parameter. To overcome this issue, before running ICA, the optimal number of components were automatically identified by GIFT toolbox.

## 5. Conclusion

The results of the present study indicate altered resting-state functional connectivity in the DMN in patients diagnosed with schizophrenia compared to matched healthy individuals. This result adds further empirical evidence for early and influential theories suggesting neural network dysconnections to account for symptoms associated with the diagnosis of schizophrenia (Andreasen et al., 1998; Friston and Frith, 1995). It also compliments previous studies reporting DMN alterations in schizophrenia. More specifically, we observed that reduced connectivity in the precuneus of the DMN was related to the severity of negative symptoms, more explicitly to the domain of apathy. In summary, the current findings emphasize the crucial role alterations in the DMN might play in schizophrenia and especially in a common negative symptom, namely apathy.

## Supporting information

