## Supplementary material for "Reduced Resting-State Connectivity in the Precuneus is correlated with Apathy in Patients with Schizophrenia"

### Methods

Participants

31 patients included in this study were receiving antipsychotic medication (amisulpride, *n*=12; risperidone, *n*=5; aripriprazole, *n*=3; quetiapine, *n*=4; clozapine, *n*=3; paliperidone, *n*=2; flupentixol, *n*=1; haloperidol, *n*=1; promethazine, *n*=1; pipamperon,*n*=2; fluphenazine, *n*=1; ziprasidone, *n*=1; chloproxiten n=1; flupenazyn n=1).

### Results

*Analysis controlling for the effects of and medication.* To assess whether medication influences the relationship between resting state connectivity and psychopathology, chlorpromazine-equivalents (CPZ) [50] were calculated. CPZ did not correlate with precuneus ($r(34) = -0.274$, $p = 0.117$). A partial correlation between the ICA parameters of the precuneus and the SANS composite score was still significant when controlling for the influence of CPZ ($r(31) = -0.367$, $p = 0.036$) as well as between the ICA parameter of the precuneus and the SANS domain of apathy ($r(31) = -0.513$, $p = 0.002$) as well as with the multimodal apathy score ($r(31) = -0.483$, $p = 0.004$)

*Analysis controlling for the effects of and medication, age, and sex.* A partial correlation controlling for the influence of CPZ, age, and sex was still significant between precuneus and SANS composite score ($r(29) = -0.438$, $p = 0.014$) and multimodal apathy score ($r(29) = -0.387$, $p = 0.031$) but only marginally between the SANS domain of apathy and precuneus ($r(29) = -0.354$, $p = 0.051$).
